# A human cortical adaptive mutual inhibition circuit underlying competition for perceptual decision and repetition suppression reversal

**DOI:** 10.1101/2023.02.22.529364

**Authors:** Teresa Sousa, Alexandre Sayal, João V. Duarte, Gabriel N. Costa, Miguel Castelo-Branco

## Abstract

A model based on inhibitory coupling has been proposed to explain perceptual oscillations. This ‘adapting reciprocal inhibition’ model postulates that it is the strength of inhibitory coupling that determines the fate of competition between percepts. Here, we used an fMRI-based adaptation technique to reveal the influence of neighboring neuronal populations, such as reciprocal inhibition, in motion-selective hMT+/V5. If reciprocal inhibition exists in this region, the following predictions should hold: 1. stimulus-driven response would not simply decrease, as predicted by simple repetition-suppression of neuronal populations, but instead increase due to the activity from adjacent populations; 2. perceptual decision involving competing representations, should reflect decreased reciprocal inhibition by adaptation; 3. neural activity for the competing percept should also later on increase upon adaptation. Our results confirm these three predictions, showing that a model of perceptual decision based on adapting reciprocal inhibition holds true. Finally, they also show that the well-known repetition suppression phenomenon can be reversed by this mechanism.

**Significance Statement:** fMRI-based adaptation has been developed as a tool to identify functional selectivity in the human brain. This is based on the notion that stimulus-selective adaptation leads to direct response suppression. In this study, we go a step further by showing that adaptation can also reveal the influence of neighboring neuronal populations. Our data reveals neural evidence for a disinhibition effect as a result of the adaptation of adjacent populations, which is in line with the adapting reciprocal inhibition model. Reciprocal inhibition can, thus, be tracked in the human brain using fMRI, adding to the understanding of human multistable perception and the neural coding of visual information. Moreover, our results also provide a mechanism for reversal of repetition suppression.

## Introduction

Models of reciprocal inhibition between neural signals in the human visual cortex have been proposed to explain perceptual decision, but experimental evidence is lacking. The classical ‘adapting reciprocal inhibition’ model of rivalry (1, 2) suggests that the strength of inhibitory coupling determines whether the system undergoes rivalrous perceptual oscillations or remains stable. Accordingly, decreases in inhibitory coupling can lead to loss of perceptual oscillations and to fusion. This model may help explain why afferent sensory information is often ambiguous, and why stabilization of perceptual interpretation may fail.

The impact of adaptation on reciprocal inhibition among visual cortex neurons has thus been postulated as one of the mechanisms accounting for perceptual bistability (2). In other words, the account that the dominance of one percept over the competing one derives from the dominant activation of a subset of neurons encoding that perceptual interpretation may only be a part of the story. The question remains whether the simultaneous suppression of those related to the opposing representations reflects reciprocal inhibitory coupling contributing to perceptual decision.

Solomon and Kohn (3) proposed an alternative model account whereby adaptation acts on two components: directly on the neuron’s receptive field and indirectly on the nonlinear gain function that includes network effects (divisive normalization). They posited that adaptation of adjacent populations results in disinhibition, explaining experimental results in which facilitation of response to the adapted stimulus was observed rather than the traditional suppression of response/fatigue. Therefore, in addition to the Lehky model (1) that postulates adaptation of inhibition this more recent alternative model also supports a disinhibition mechanism. Here, we aimed to investigate for a neural signature of such disinhibition mechanism as probed by experimentally induced adaptation.

While revisiting Levelt’s propositions on visual perception, Brascamp et al. (4) discussed two types of perceptual alternation: the escape and the liberation transitions. In both cases, the perceptual shift is suggested to be caused by an increase in the activity of the inhibited populations rather than a decrease in the activity of the dominant population. In the first case, the activity of the suppressed neuronal population increases to a critical level. In the second, the activity of the dominant population decreases, by adaptation, to a critical level. The recovery of the suppressed population activity is attributed to a recovery of that population’s adaptation (an exit from the fatigue/hyperpolarization state), rather than a reduction of inhibition of the dominant population.

Functional magnetic resonance imaging (fMRI)-based adaptation has consistently been shown to reflect stimulus selectivity by response reduction, defining repetition suppression. This technique became a popular method to show selectivity of populations of neurons. Its validity was demonstrated by comparing single-cell recordings with functional imaging of orientation, motion, and face processing (5), which showed remarkable consistency across experimental models. However, positive phenomena related to increases in neural activity, remain unknown or even disputed- such as the prior claim that motion aftereffects lead to an increase in activity in the motion-selective hMT+/V5 (6). Accordingly, Huk et al. (7) showed that motion aftereffects do not lead to increased overall neural activation, if attention is controlled for. In the macaque middle temporal area (MT) and its human homologue (hMT+/V5), adaptation to moving stimuli reduces brain activity (8–10) supporting repetition suppression mechanisms in this region. These results fit with a view of adaptation as a passive process of neuronal fatigue: the more a neuron fires, the more its subsequent response is reduced (8).

Here, we went a step further by hypothesizing that adaptation can also reveal reciprocal inhibition in the human brain, and in particular in hMT+/V5, which may possibly revert repetition suppression. This hypothesis is based on the notion that motion-sensitive neurons in this region are not isolated feature detectors, and that populations of neurons with different preferred directions interact at a net level through reciprocal inhibition. In this case, the adaptation of one particular population of neurons can reduce the inhibitory input to other populations preferring different directions and thereby increase their response, leading to perceptual bistability, which would be measurable with fMRI.

It is important to point out that testing conflicting opposite local directions often leads to response reduction but not perceptual oscillations (a distinct phenomenon known as motion opponency). On average only minor changes are observed at the single cell level, reflected by pooling of local motion signals (5) and leading to reduced flicker processing (11). In human area MT+/V5, this is echoed by the observation that stimuli with perfectly balanced motion energy constructed from local dots moving in counter-phase elicit a weaker response than non-balanced (in-phase) motion stimuli (12).

In this study, we deal with global motion representations competing for perceptual decision – multistable perception (13–17). Competing percepts are often mutually exclusive: only one interpretation is visible at a time, while the others are suppressed from perceptual awareness (2). Current perceptual decision models consider multistability as the result of the integration of the topdown influence of high-level cognitive processes *vs*. bottom-up sensory processes (15, 18–23). Adaptation and noise are known relevant low-level components for determining perceptual multistability (24–28). However, reciprocal inhibition as revealed by adaptation has not been tested so far.

Separate pools of neurons, each eliciting one of the possible perceptual interpretations, may exert direct inhibition over each other. The model explaining why reciprocal inhibition may be key in shaping perceptual transitions in binocular rivalry by Lehky (1) allows the neuronal pool underlying one perceptual interpretation to directly suppress activity in the neuronal pool representing the other interpretation. This may help explain the timing of perceptual dominance of each of the alternatives. Neuronal adaptation by progressively weakening the dominant population causes disinhibition of the suppressed population, which might lead to multiple cycles of alternating percepts. Finally, various sources of noise contribute to the variability in the time elapsed between successive transitions (27, 29, 30) as also implemented in Lehky’s model.

Data from psychophysics and brain imaging need to be anchored on computational models to explain bistability (31–38). We tested the adapting reciprocal inhibition model in our neurobehavioral approach. In a pure and direct adaptation model (without reciprocal interactions) the activation strength should weaken throughout a dominance period as adaptation weakens the dominant response, and it should be weakest just before a switch in dominance (2). If adaptation is combined with reciprocal inhibition this would imply a faster switching mechanism, because the dominated representation would be released from inhibition due to adaptation of the dominant population, and increase its activity faster. We thus tested, using functional neuroimaging, for signatures of reciprocal inhibition as probed by adaptation.

Our experimental design was partly inspired by behavioral studies showing the ‘‘reverse-bias effect”, a phenomenon in which a prolonged presentation of an unambiguous stimulus leads subjects to report the opposite interpretation during a subsequent presentation of the ambiguous stimulus (28). Exposure to the unambiguous figure presumably adapts the neuronal pool supporting the corresponding percept, so that when the ambiguous figure is presented, these structures cannot compete effectively with the opposing neuronal pool, and the opposite interpretation of the stimulus is perceived. However, the question remains whether pure adaptation per se is sufficient to explain the suppression of the adapted percept or if reciprocal inhibition between neuronal populations also plays a role. This question also remains open in experiments with ambiguous stimuli (28) and binocular rivalry (34).

Following this question, we tested whether reciprocal inhibition mechanisms contributing to perceptual bistability can be revealed at the neural level by using adaptation as a probe. We made the following predictions: 1. Reciprocal inhibition should be revealed by fMRI adaptation, manifested as increases of activity from adjacent populations leading to increased net activity, instead of the decreases predicted by simple repetition suppression; 2. In perceptual decision tasks involving competing motion representations, perceptual preference should reflect the effect of adaptation on reciprocal inhibition; 3. Neural activity for the competing perceptual representation should also later on increase upon adaptation (a neural “reverse bias” effect). This framework, highlighted in Figure 1, predicts that neuronal adaptation of one surface representation leads to neuronal enhancement of a competing one, in addition to what would be expected by pure adaptation alone to the same stimulus.

**Figure 1.**
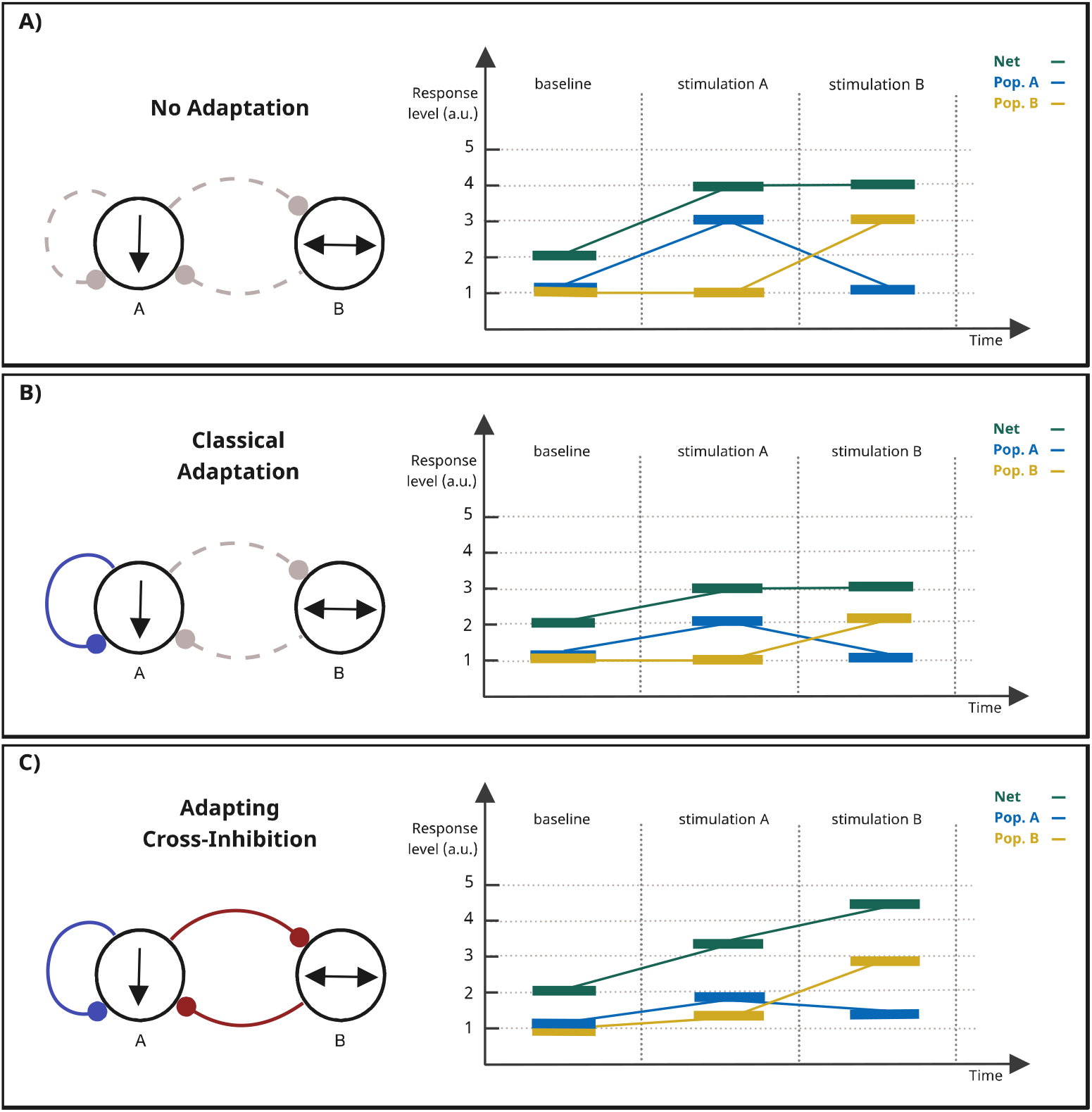
Predictions generated by the adapting reciprocal inhibition model at the neural level. This model explains why adaptation may surprisingly lead to increased neural net response and changes in perceptual interpretation. A) Expected neural response patterns in the absence of adaptation. B) Expected neural response patterns in the presence of adaptation. C) Expected neural response patterns in the presence of adapting cross-inhibition. Population A (Pop. A) is here representing the neural population being first directly stimulated (stimulation A/adapting period). Population B (Pop.B) comprises the full set of neighboring populations including (but not exclusively) the one that is sensitive to stimulation B. In other words, a large set of populations can be disinhibited by adapting cross-inhibition, contributing to an overall increase in activity that overcomes the initial decrease predicted by classical adaptation. Gray lines: inactive mechanisms; Blue lines: active adaptation, Red Lines: active adaptive cross-inhibition.

As a model region to study this problem, we used the hMT+/V5 brain region, which is an early-level node in perceptual decision regarding integration or segregation of surface motion (23, 39). Early work in the awake behaving macaque linked responses in the functional domains of extrastriate area MT with perception of motion, which was later extended to humans (6, 35, 40–42). We applied a well-studied ambiguous stimulus paradigm, the moving plaids (Figure 2), that elicit perceptual bistability (43, 44). This paradigm uses different moving surfaces that can be perceptually integrated into two alternative ways, corresponding to the activation of distinct neuronal populations. It does therefore provide a critical opportunity to study mechanisms of interaction between the neuronal pools in human motion complex representing different perceptual domains and define, based on adaptation, how they influence the final percept.

**Figure 2.**
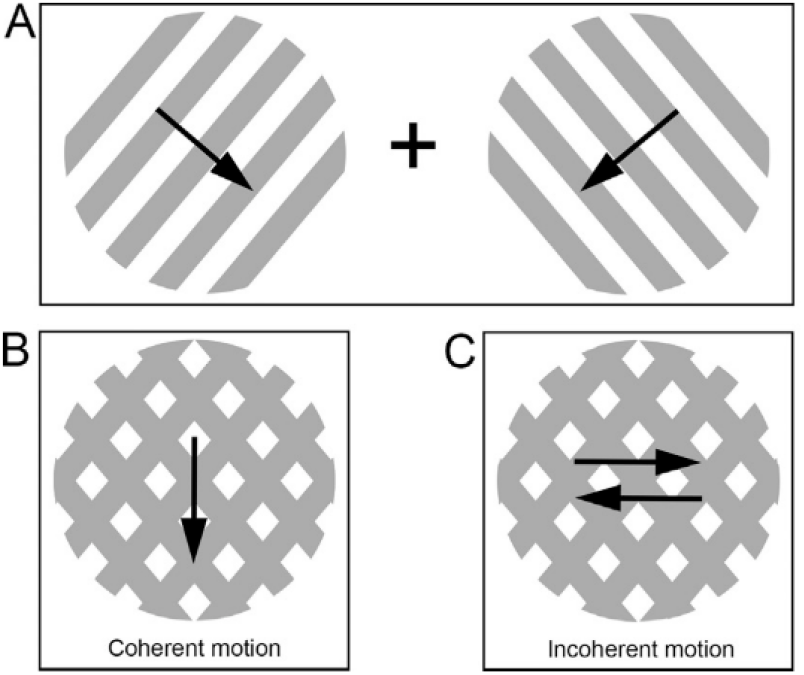
Plaid stimulus illustration. By superimposing two gratings (A), moving orthogonally to the gratings’ orientation, a moving ambiguous stimulus is created. Moving plaid stimuli elicit two alternative perceptual interpretations and allow for the study of bistable perceptual transitions between integration (one surface, coherent motion - B) and segmentation (two surfaces, incoherent motion - C) of the visual cues. Arrows depict the number and direction of perceived moving surfaces.

Previously, we have found evidence for distinct levels of adaptation during both percepts of this type of stimulus (45) and that adaptation competes with short-term visual memory mechanisms for perceptual interpretation (46). Here, we tested for evidence of an adapting reciprocal inhibition mechanism in the same brain region. This was, here, done by measuring hMT+/V5 blood-oxygen-level-dependent (BOLD) responses for each of the two alternative percepts, while imposing long adaptation periods to either one of the percepts.

## Results

### Behavioral results

We started by evaluating how perceptual adaptation influenced perceptual dominance during ambiguous visual stimulation. Statistical analysis shows that perception depended on the previously adapting representation (Figure 3). During the ambiguous period after adapting to the coherent percept, participants mainly reported seeing incoherent motion, while after adapting to the incoherent percept, participants mainly reported seeing coherent motion.

**Figure 3.**
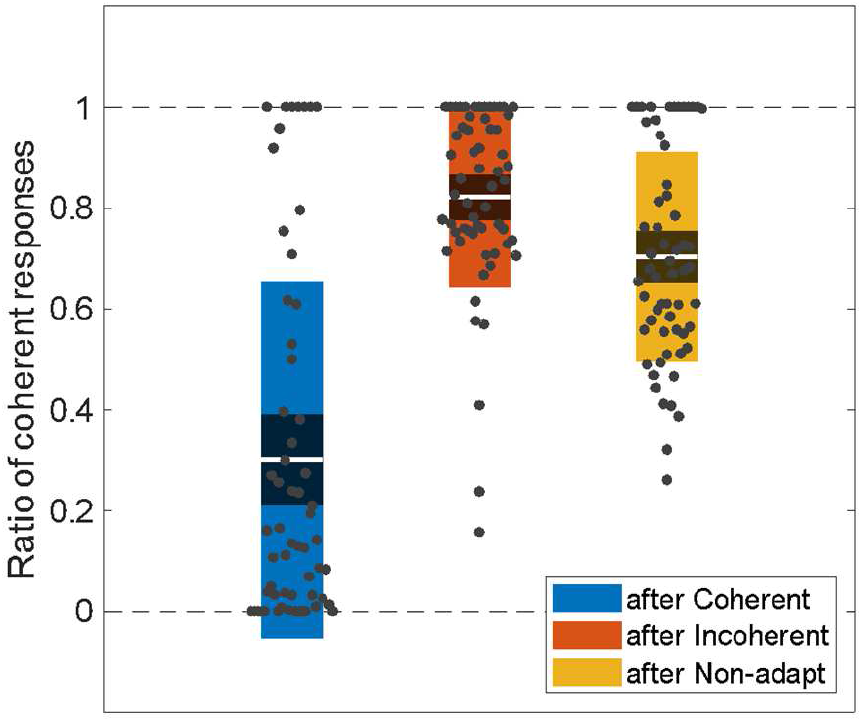
Perceptual dominance depended on previous adaptation to different percepts (*test of prediction 2 -* reverse bias effect). We display the ratio of coherent percepts during plaid ambiguous motion depending on the previous adapting/non-adapting period (after coherent, incoherent, or non-adapting). Each data point corresponds to a single run and is displayed over the 95% confidence interval (dark color area) and one standard deviation (light color area). The mean of the values is represented as a white line.

A repeated measures one-way ANOVA revealed a significant main effect *F*(2, 19) = 41.4, (*p* = 1.30×10^-6^) of the adapting condition on the perceptual dominance during plaid ambiguous motion. A post-hoc Tukey’s multiple comparisons test specifically showed that the mean ratio of coherent responses in the ambiguous period was significantly higher (*p* = 8.56×10^-6^) after adapting to incoherent (0.82 ± 0.17) than after adapting to coherent motion (0.30 ± 0.35). Moreover, the differences between both adapting and the non-adapting trials were also significant (‘after coherent *vs*. non-adapting’: *p* = 6.82×10^-6^; ‘after incoherent *vs*. non-adapting’: *p* = 1.14×10^-3^). Overall these results point to a reverse bias effect, and therefore for a behavioral signature of reciprocal inhibition, consistent with prediction 2.

### fMRI results

Neurophysiological data were first used to localize the hMT+/V5 complex of each participant. Based on the contrast between moving and static conditions of the localizer run, we were able to functionally define this region on the left and right hemispheres. The group-average MNI coordinates (*x, y, z*) for the left hMT+/V5 are (−47,-72,6) and for the right hMT+/V5 are (47,-69,3). We then analyzed the hMT+/V5 BOLD response during the entire trial, as shown in Figure 4.

**Figure 4.**
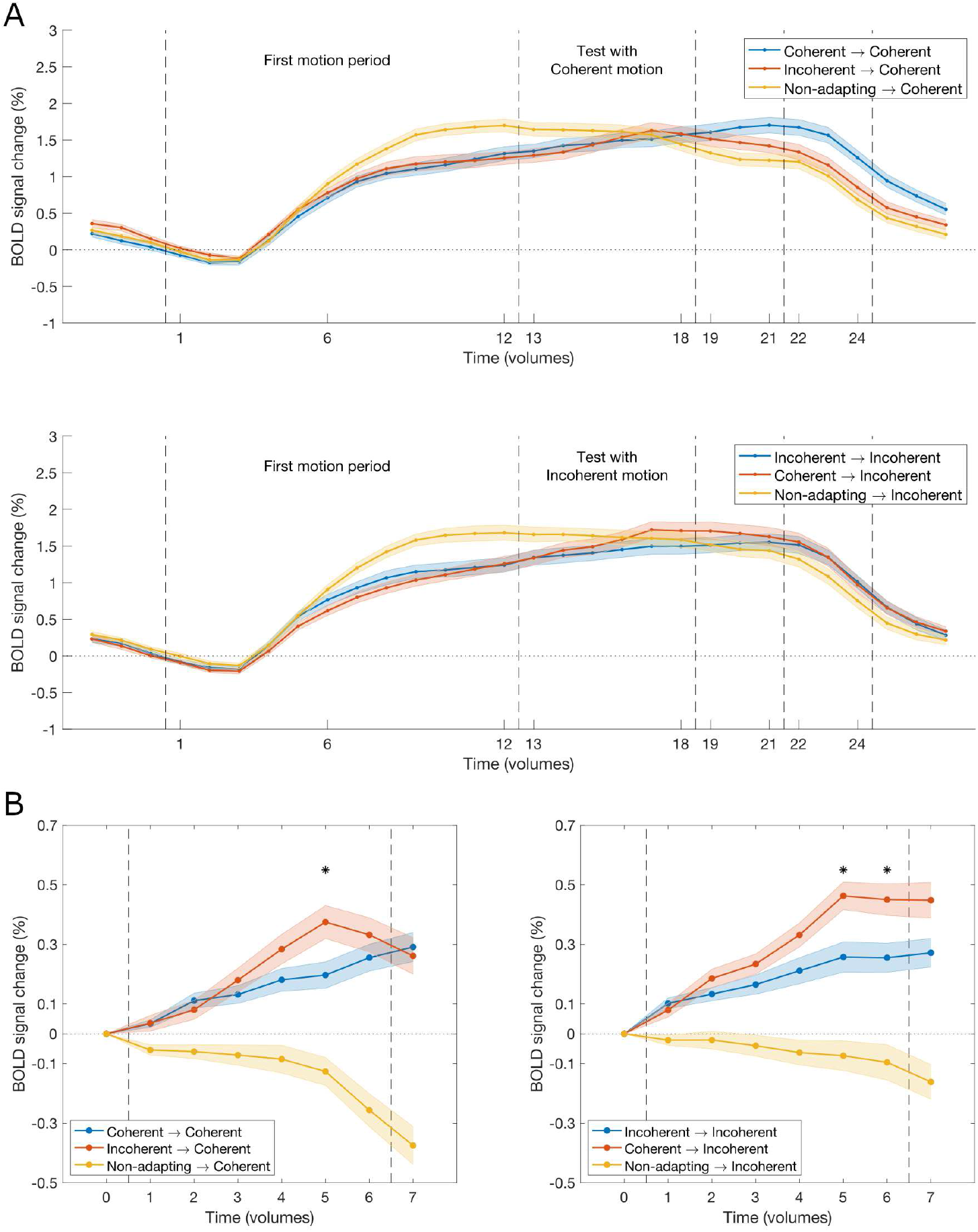
hMT+/V5 response during adapting/non-adapting and testing conditions. (A) Test of prediction 1: in spite of the expected initial difference between adapted and non-adapted conditions, there is a subsequent increase in brain activity already during the adaptation period (first motion period). We display the bilateral hMT+/V5 percent signal change during the entire trial relative to the *Static* condition. Responses to coherent (top) and incoherent (bottom) percepts are shown after adaptation to coherent motion, after adaptation to incoherent motion, and after non-adapting motion. (B) Test of prediction 3: neural activity for coherent (left) and incoherent (right) motion increases upon adaptation to the competing percept. The curves were normalized to start from zero. Statistical significant differences between the adapting curves at each time point are marked (*) when significant between all conditions (*p* < 0.05, corrected for multiple comparisons). In both panels, the shaded area displays the standard error of the mean.

During the first motion period, the BOLD signal in hMT+/V5 increased differently for the three types of motion. Contrasting with the response to the non-adapting motion, the response to both adapting stimuli was initially smaller, as predicted by conventional adaptation and repetition suppression models but then showed a gradual and surprising increase in activity throughout the block, in agreement with our prediction number 1.

In the second motion period, the signal in hMT+/V5 also varied depending on the trial. Here, we highlight the trials where opposing percepts were shown (Figure 4, red lines): in these trials, BOLD activity peaked even further after the transition to the opposing percept, suggesting disinhibition due to lack of suppression of the adapted population, according to our prediction number 3.

The hMT+/V5 responses to the coherent and incoherent plaid motion were compared at each volume of the testing period between the three conditions. During the test to coherent plaid motion (Figure 4B, left panel), the normalized signal was significantly higher after adapting to the incoherent percept than after adapting to the coherent one (*p* = 2.10×10^-2^, corrected) and also after the non-adapting period (*p* = 1.49×10^-5^, corrected) 5 volumes (= seconds) after the switch.

During the incoherent plaid motion test (Figure 4B, right panel), the hMT+/V5 response was significantly higher after adapting to the coherent percept than after adapting to the incoherent one (*p* = 1.22×10^-2^, corrected) and also after the non-adapting period (*p* = 3.26×10^-4^, corrected) 5 and 6 volumes (= seconds) after the switch.

An important remark of this analysis is that the hMT+/V5 signal tended to decrease continuously after the non-adapting period, but to increase after previous adaptation to the same or opposing percept, further reinforcing prediction 3. This was confirmed by a linear regression analysis comparing ‘coherent after incoherent’ (slope = 0.05, *R^2^* = 0.78) and ‘coherent after non-adapting’ (slope = −0.05, *R^2^* = 0.81) revealing a significant difference between them (*F*(1,12) = 46.44, *p* = 1.86×10^-5^). The same is true when comparing ‘incoherent after coherent’ (slope = 0.07, *R^2^* = 0.94) and ‘incoherent after non-adapting’ (slope = −0.02, *R^2^* = 0.90) revealing a significant difference between them (*F*(1,12) = 129.10, *p* = 8.87×10^-8^).

## Discussion

In this study, we tested the adapting reciprocal inhibition hypothesis at the perceptual decision and neural level. This hypothesis was based on three predictions:

### Prediction 1

Adaptation should decrease inhibition from the dominant perceptual representation and release adjacent neuronal populations, with a direct impact on neural responses to the opposite representation, which we indeed found to be increased. Contrary to the expected decrease due to simple repetition suppression, adaptation curves showed a gradual increase, providing further support to our hypothesis.

The hMT+/V5 responses were initially lower than during non-adapting conditions and this effect was stronger during the adaptation to coherent motion pattern (the expected difference induced by adaptation). Neuronal adaptation, also known as repetition suppression, is known to result from repeated stimulation and to lead to a smaller amplitude of the measured neurophysiological signal (5, 47, 48). In a previous study, we found that during moving plaid visualization, visual adaptation was stronger upon coherent and into a lesser extent, incoherent percepts (45).

However, the hMT+/V5 response to both adapting stimuli gradually increased throughout the entire adaptation period, challenging the expected activity decrease by simple repetition suppression accounts (49). This is evident during the adaptation blocks and extends to the tests where the motion patterns remain the same, extending the adaptation period (Figure 4). An explanation, in line in particular with the models proposed by Lehky (1) as well as by Solomon and Kohn (3), for the observed late increase of hMT+/V5 net activity is that it reflects the weakening of the suppression strength of one perceptual domain over the other. This allows for the increase of activity of the latter (which would not occur without the adaptation effect on reciprocal inhibition). Such finding represents functional neuroimaging evidence for the major prediction of the adapting reciprocal inhibition model (1, 2). It is consistent with sustained adaptation of the stimulated perceptually selective population and the consequent gradual disinhibition of the adjacent populations. Additionally, the time-course of hMT+/V5 response to prolonged coherent motion tends to become higher than the response to the incoherent motion, which also matches the hypothetical disinhibition effect (stronger for the largest adapting condition). In other words, taking into account the stronger neuronal adaptation to this motion pattern (45), the disinhibition effect is also expected to be stronger.

### Prediction 2

A correlate of the adaptation effect on reciprocal inhibition mechanisms should also be evident at the perceptual decision level. Accordingly, we found that ambiguous moving plaid interpretation depended on previous adaptation to the alternative percept (Figure 3). This is consistent with a reverse-bias effect, as a signature of reciprocal inhibition. Such a phenomenon had also been suggested to occur in binocular rivalry (34).

### Prediction 3

The adaptation of neuronal populations sensitive to the coherent plaid motion should increase the net neural response to a subsequently presented incoherent plaid motion version, which activates opposing populations, and vice versa. Indeed, hMT+/V5 activity peaked after transitioning to the opposing percept. This finding further corroborates the notion that reciprocal inhibition varies over time as a function of adaptation, which affects neighboring neuronal populations tuned to distinct global motion representations. It also provides evidence for a low-level neural mechanism underlying the interaction between the competing perceptual states.

hMT+/V5 contains columnar populations, each column of neurons representing a global direction of motion. Columns preferring a given direction provide local cross-inhibitory influences to columns preferring different directions while providing long-range excitation to columns responding preferentially to the same direction (9, 50). In line with that, our results are consistent with the hypothesis that, on the one hand, motion adaptation in a given direction leads to local adaptation of the columns responding to that direction; on the other hand, they are consistent with the hypothesis that adaptation releases neighboring columns from inhibition, thereby allowing their respective perceptual representation to eventually win and provide a switch in the perceptual decision.

In sum, this study provides neural evidence for a reciprocal inhibition model and architecture in human visual cortex with a direct impact on perceptual decision (1, 3, 34). Our data support the hypothesis of adaptive inhibition, which significantly influences neural activity and perception. Importantly, the link between these both neuronal mechanisms may shed light on phenomena such as the reverse perceptual bias. Finally, our work shows, for the first time, that fMRI BOLD data can reveal time-varying disinhibition as a function of neuronal adaptation, and evidence for reversal of repetition suppression mechanisms.

## Materials and Methods

### Participants

20 volunteers were recruited to participate in this study (9 male, mean age 27.7 ± 4.6 years). All had normal or corrected-to-normal vision and no history of neurological or psychiatric diseases, and were right-handed (mean laterality index of 80.3 ± 17.8) (51). All gave informed written consent, in accordance with the declaration of Helsinki, and the study complied with the safety guidelines for magnetic resonance imaging research on humans. The study was approved by the Ethics Committee of the Faculty of Medicine of the University of Coimbra.

### Stimulus

We used a well-known ambiguous stimulus - moving plaids - which allows the investigation of switches between integration and segmentation of motion vectors (43, 44, 52). Plaid stimuli lead to bistable interpretation because of spontaneous switches between perceptual integration (coherent motion) and segregation (incoherent motion) of the visual cues (Figure 2).

Here, to induce the perception of each one of the two possible motion interpretations, we added dots to the plaid (superimposed on the bars) to disambiguate motion (Supplementary Figure 1). Depending on the relative percentage of dots that move vertically or horizontally, we were able to induce a given level of coherence, hence a perceptual interpretation to the participant. With 100% of the dots moving vertically (down), the coherent percept is induced, while with 100% of the dots moving horizontally (half to the left and half to the right), the incoherent percept is induced. The ambiguous version of the plaids was obtained using the same plaid version without any added dots. A complete description of the stimulus properties is provided in Supplementary Table 1, matching our previous studies (45, 46). We used a total of four different configurations of the stimulus to induce different perceptual outputs: unambiguous coherent motion (all the dots moving down), unambiguous incoherent motion (half the dots moving left and the other half moving right), alternating between coherent and incoherent motion (referred here as non-adapting motion - the motion direction and coherency changes at every time sample) and finally ambiguous motion (without any dots).

Visual stimuli were created in MATLAB R2016b (The Mathworks, Inc., Natick, MA-USA) running Psychophysics Toolbox version 3 (53, 54).

### Experimental design

This study was organized into two experiments. The first one consisted of a behavioral test outside of the magnetic resonance imaging (MRI) scanner. It aimed to test whether the perceptual report during ambiguous stimulation depended on a previous adaptation period, as evidence for the reverse-bias effect on the perception of the moving ambiguous plaid. The second one consisted of an fMRI session that allowed us to explore the neural correlates of cross-inhibitory mechanisms underlying bistable perception, in the same participants.

#### Behavioral testing

We evaluated whether the adaptation to an unambiguous version of the plaid stimulus leads to a predominant percept during a subsequent ambiguous version of the moving plaid. The session lasted approximately 16 minutes, comprising three runs. Each run consisted of 12 stimulation trials (Supplementary Figure 2) including: an initial static interval (non-moving plaid with dots) of 6 s; an adapting or non-adapting motion condition (constantly coherent or incoherent motion leading to adaptation, and rapid alternation between both avoiding adaptation effects, respectively) of 12 s; followed by the ambiguous version of the moving plaid for 6 s, with a final short no motion interval of 3 s. During the ambiguous motion condition (the second motion block), participants were asked to look at the central fixation cross and to continuously report, via button press, which percept (coherent or incoherent) they were experiencing. The last no motion block was used to encompass the expected motion after-effect (MAE) and allow it to fade out before restarting the next trial with the static plaid. The 12 stimulation trials consisted of four repetitions of the test for each type of adaptation – coherent or incoherent motion – and four repetitions of the test for the control/non-adapting stimulation. The sequence of trials was randomized between runs and participants.

Stimuli were presented on a computer monitor (resolution: 1920 x 1080 pixels, refresh-rate: 60 Hz) at 70 cm from participants. Button presses were recorded using a standard mechanical keyboard at 60 Hz.

#### fMRI study

We recorded the hMT+/V5 response to the coherent and incoherent plaid motion after adapting to the coherent and incoherent plaid motion, and after a non-adapting condition, using fMRI. As we aimed at studying the neuronal responses to each percept after adapting to the opposing one, the ambiguous period (used in the behavioral experiment) was replaced by either coherent or incoherent motion (Supplementary Figure 3). By doing this, we ensured the same number of tests per percept and avoided perceptual switches during the test period, while testing for the effects of adaptation on the opposite percept, to investigate signatures of cross representation interactions.

Participants’ attention was controlled through a counting task related to the central fixation cross. It allowed us to control for adaptation bias due to shifts in arousal or by selective attention, as previously described by Castelo-Branco *et al* (55). From one to four times in each trial, the central fixation cross became slightly larger (0.67° to 0.80°) for 250 ms. The task was to detect and count these changes, and to report this number at the end of the trial, as previously done by Sousa *et al*. (45).

Each trial included a static condition (static plaid stimulus with superimposed dots) of 6 s, followed by an adapting or non-adapting plaid moving condition of 12 s, a test condition of 6 s, a second no motion period of 3 s, and a report period of 3 s. The second no motion block was used to encompass the expected MAE. During the report period, the participant used one of four buttons of a response box to report the number of cross size changes (1 to 4 changes). Within the report block, feedback (correct/incorrect) was provided by briefly changing the color of the fixation cross to green or red, respectively.

In each run, we included 12 trials, i.e.- two trials of each type: coherent after coherent, coherent after incoherent, coherent after non-adapting, incoherent after incoherent, incoherent after coherent, and incoherent after non-adapting. The sequence of trials was randomized between runs and participants. Participants performed 6 runs, in a total of 45 minutes.

### Behavioral data analysis

The perceptual reports acquired during the ambiguous stimulus were analyzed by evaluating the number of each type of report (in number of frames), coherent and incoherent, per trial, and its average duration. Then, the ratio of coherent responses, calculated as the number of ‘coherent’ reports over the total number of reports during the ambiguous period, was estimated for each trial type – after coherent, incoherent- and non-adapting motion.

A repeated measures one-way ANOVA, with Greenhouse-Geisser correction as implemented in IBM (Armank, 169 NY) SPSS Statistics 22.0 software package, was applied to test for the presence of a main effect of the trial type on perceptual reports. Differences between the three types of trials were tested based on post-hoc Tukey’s multiple comparisons test.

### fMRI data acquisition

Scanning was performed on a 3T Siemens Magnetom Prisma Fit scanner at the Institute of Nuclear Sciences Applied to Health, Portugal, using a 64-channel head coil. The scanning session started with the acquisition of one 3D anatomical magnetization-prepared rapid acquisition gradient echo (MPRAGE) pulse sequence (TR = 2530 ms, echo time (TE) = 3.42 ms, flip angle = 7°, 176 slices, voxel size 1.0 × 1.0 × 1.0 mm^3^, field of view (FOV) = 256 × 256 mm). Then, six functional runs were acquired using a multi-band accelerated echo planar imaging (MB-EPI) sequence from the Center for Magnetic Resonance Research, University of Minnesota (Release R016a). The main parameters were: multi-band factor = 3, TR = 1000 ms, TE = 30.2 ms, flip angle = 68°, 42 interleaved slices with 0.5 mm gap, voxel size 2.5 × 2.5 × 2.5 mm, FOV 192 × 192 mm, number of volumes = 374. Cardiac pulse signal was acquired during the functional runs, using the proprietary peripheral pulse unit from Siemens. In total, the session lasted for approximately 45 minutes.

During the fMRI session the stimulus was presented on an LCD screen (70 x 39.5 cm, 1920 x 1080 pixel resolution, 60 Hz refresh rate) which the participants viewed through a mirror mounted above their eyes at an effective distance of 156 cm. Participants’ reports were recorded using a fiber-optical MR-compatible response box (Cedrus Lumina LSC-400B). To confirm whether participants maintained central fixation during the acquisition session, individually calibrated eye-tracking data (sample frequency of 500 Hz) were recorded inside the scanner using Eyelink 1000 software (SR Research, Ottawa, Canada).

### fMRI data processing

Preprocessing was performed using SPM12 (Wellcome Department of Cognitive Neurology, London; http://www.fil.ion.ucl.ac.uk/spm) running on MATLAB 2018b (The MathWorks Inc., Natick, MA). Functional images were slice-time corrected, realigned, normalized to the Montreal Neurological Institute (MNI) template space, and spatially smoothed with a 5 mm full width at half maximum Gaussian kernel.

After preprocessing, all data were modeled with a general linear model (GLM), accounting for autocorrelations with FAST. The design matrix included a high-pass filter with a cutoff period of 60 s and regressors for all stimulation blocks convolved by the canonical hemodynamic response function. Based on the cardiac signal recorded during the session, the cardiac phase time series were generated using the PhysIO toolbox (Kasper *et al*., 2017a) implementation for the RETROICOR method. Four cardiac nuisance regressors were added to the design matrix, together with the six head motion parameters.

### fMRI data analysis

#### hMT+/V5 localization

The left and right hMT+/V5 were functionally localized in every participant with an individual GLM analysis of the first functional run. The regions were defined as a 5 mm sphere centered at the voxel in the posterior middle temporal region responding most significantly to the balanced contrast “Adapt coherent + Adapt incoherent + Non-adapt > Static”. The individual statistical maps were limited at threshold value of *p* = 0.01, *FWE-corrected*. For the following analysis, the left and right ROIs were used combined as a single region, which we designate by bilateral hMT+/V5.

#### hMT+/V5 response analysis

The bilateral hMT+/V5 ROI time courses were extracted for each run and participant, and then normalized as percent signal variation, using as baseline the average value during the “Static” condition. These normalized time courses were split into trial type and averaged across trials/runs and participants, resulting in a group analysis of the BOLD activation in bilateral hMT+/V5.

To further analyze the BOLD signal dynamics during the second motion period, and its modulation dependence on the preceding adapting condition, we have compensated for the different initial level of activity (at the beginning of this period) for each of the three trial types, by subtracting the average value of the three data points before the period of each trial type. In this way, we ensured that the time course differences did not depend on the initial signal value, but rather on the stronger or weaker level of activity in hMT+/V5. A repeated measures one-way ANOVA, with Greenhouse-Geisser correction as implemented in SPSS, was applied at each time point to test for differences between all types of trial. Post-hoc comparisons were run based on post-hoc Tukey’s test. Moreover, a regression model was used to determine whether there was a significant difference between the increase/decrease response recorded after adapting to the opposing percept and after non-adapting conditions.

## Code availability

The MATLAB code used for the stimulus and analyses performed in this study can be found at https://github.com/CIBIT-UC/public_vpinhibition.

## Acknowledgments

We would like to thank the participants for their involvement in this study. We are also very grateful to Sónia Afonso and Tânia Lopes for the help with fMRI setup and scanning, and to Ricardo Martins for the valuable discussions about stimuli design and data analysis.

