## Supplementary information for "A human cortical adaptive mutual inhibition circuit underlying competition for perceptual decision and repetition suppression reversal"

**This PDF file includes:**

Figures S1 to S3
Tables S1

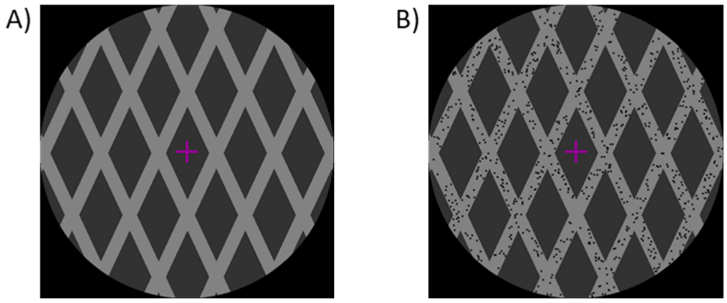


**Fig. S1.** Stimuli used in the experiment. To force the perception of one of the two possible motion interpretations (coherent and incoherent motion), we added dots to the ambiguous moving plaid (A) to disambiguate motion (B). Depending on the moving dots' direction (superimposed on the bars), the otherwise ambiguous stimulus was readily perceived as a plaid moving coherently or incoherently.

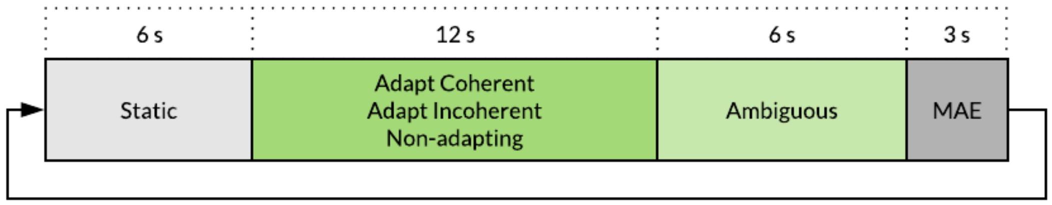


**Fig. S2.** Experimental protocol diagram for behavioral session. Adaptation of 12 seconds long was used to test the reverse-bias effect on the perception of the moving ambiguous plaid, which is believed to highlight cross-inhibitory mechanisms. Each stimulation trial started with a static version of the unambiguous plaid, followed by the adaptation to coherent or incoherent motion and the test with ambiguous motion, which finished with a short no motion period to avoid motion after-effect (MAE) bias on the subsequent trial. A non-adapting condition was used as a control (for details see Methods and Supplementary Data).

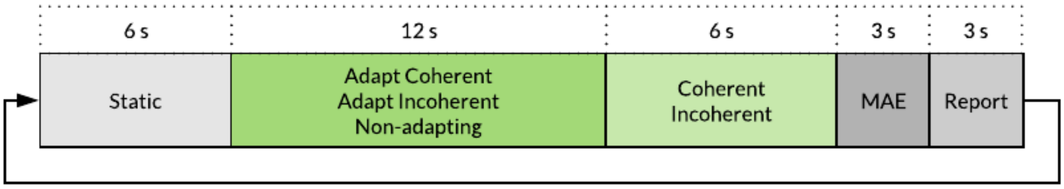


**Fig. S3.** Experimental protocol diagram for fMRI session. Twelve seconds long adaptation periods were used to test the hMT+/V5 response to the incoherent and coherent motion plaid after perceptual adaptation to the opposing motion pattern. A non-adapting condition was used as a control for the adaptation effects. To avoid data bias due to attentional shifts participants were asked to count the number of times that the central fixation cross varied its size during the trial and to report it at the end.

**Table S1.** Plaid stimulus properties.

| | |
|---|---|
| Angle of gratings relative to horizontal (º) | 65 |
| Duty cycle (%) | 25 |
| Aperture diameter (º visual angle) | 9 |
| Screen background color (RGB) | (0, 0, 0) |
| Plaid background color (RGB) | (50, 50, 50) |
| Gratings color (RGB) | (130, 130, 130) |
| Spatial frequency (cycle/º visual angle) | 0.625 |
| Motion speed (º visual angle/s) | 1.6 |
| Number of dots | 800 |
| Dots color (RGB) | (20, 20, 20) |
| Dots size (º visual angle) | 0.05 |
| Dots horizontal speed (º visual angle/s) | 2.4 |
| Dots vertical speed (º visual angle/s) | 4 |
| Fixation cross width (º visual angle) | 0.67 |
